# An integrative strategy to identify the entire protein coding potential of prokaryotic genomes by proteogenomics

**DOI:** 10.1101/153213

**Authors:** Ulrich Omasits, Adithi R. Varadarajan, Michael Schmid, Sandra Goetze, Damianos Melidis, Marc Bourqui, Olga Nikolayeva, Maxime Québatte, Andrea Patrignani, Christoph Dehio, Juerg E. Frey, Mark D. Robinson, Bernd Wollscheid, Christian H. Ahrens

## Abstract

Accurate annotation of all protein-coding sequences (CDSs) is an essential prerequisite to fully exploit the rapidly growing repertoire of completely sequenced prokaryotic genomes. However, large discrepancies among the number of CDSs annotated by different resources, missed functional short open reading frames (sORFs), and overprediction of spurious ORFs represent serious limitations.

Our strategy towards accurate and complete genome annotation consolidates CDSs from multiple reference annotation resources, *ab initio* gene prediction algorithms and *in silico* ORFs in an integrated proteogenomics database (iPtgxDB) that covers the entire protein-coding potential of a prokaryotic genome. By extending the PeptideClassifier concept of unambiguous peptides for prokaryotes, close to 95% of the identifiable peptides imply one distinct protein, largely simplifying downstream analysis. Searching a comprehensive *Bartonella henselae* proteomics dataset against such an iPtgxDB allowed us to unambiguously identify novel ORFs uniquely predicted by each resource, including lipoproteins, differentially expressed and membrane-localized proteins, novel start sites and wrongly annotated pseudogenes. Most novelties were confirmed by targeted, parallel reaction monitoring mass spectrometry, including unique ORFs and variants identified in a re-sequenced laboratory strain that are not present in its reference genome. We demonstrate the general applicability of our strategy for genomes with varying GC content and distinct taxonomic origin, and release iPtgxDBs for *B. henselae*, *Bradyrhozibium diazoefficiens* and *Escherichia coli* as well as the software to generate such proteogenomics search databases for any prokaryote.

## Introduction

Advances in next generation sequencing technology and genome assembly algorithms have fueled an exponential growth of completely sequenced genomes, the large majority of which (>90%) originate from prokaryotes (Reddy et al. 2015). The accurate annotation of all protein-coding genes (interchangeably used with CDSs from here on) is essential to exploit this genomic information at multiple levels: from small, focused experiments, up to systems biology studies, functional screens and accurate prediction of regulatory networks.

Yet, obtaining a high quality genome annotation is a challenging objective. Pipelines for automated *de novo* annotation of prokaryotic genomes have been developed (Aziz et al. 2008; Markowitz et al. 2009; Davidsen et al. 2010; Vallenet et al. 2013). Such annotations greatly benefit from a manual curation step to catch obvious errors (Richardson and Watson 2012), which is carried out for selected reference genomes by resources like NCBI’s RefSeq (Pruitt et al. 2012) or Microscope (Vallenet et al. 2013). Major re-annotation efforts can affect hundreds of CDSs (Luo et al. 2009), highlighting the relevance of accurate genome annotations (Petty 2010).

Despite improvements in functional genome annotation, three major issues remain: the discrepancies of the number of CDSs annotated by different reference annotation resources (Poole et al. 2005; Bakke et al. 2009; Cuklina et al. 2016), the over-prediction of spurious ORFs that do not encode a functional gene product (Dinger et al. 2008; Marcellin et al. 2013), and the underrepresentation of short ORFs (sORFs) (Hemm et al. 2008; Warren et al. 2010; Storz et al. 2014). True sORFs, which often belong to important functional classes like chaperonins, ribosomal proteins, proteolipids, stress proteins and transcriptional regulators (Basrai et al. 1997; Zuber 2001; Hemm et al. 2008), are inherently difficult to differentiate from the large amount of spurious sORFs (Dinger et al. 2008; Marcellin et al. 2013).

Proteogenomics, a research field at the interface of proteomics and genomics (Nesvizhskii 2014), is one attractive approach to address these problems. The direct protein expression evidence provided by tandem mass spectrometry (MS) for CDSs missed in genome annotations (proteogenomics) differs from ribosome profiling data: while the latter can capture translational activity on a genome-wide scale (Ingolia 2014), the former allows detection of stable proteins. First used in the genome annotation effort for *Mycoplasma mobile* (Jaffe et al. 2004), proteogenomics has since been applied to both prokaryotes (Gupta et al. 2007; de Groot et al. 2009; Payne et al. 2010; Venter et al. 2011; Kumar et al. 2013; Marcellin et al. 2013; Kucharova and Wiker 2014; Cuklina et al. 2016) and eukaryotes (Nesvizhskii 2014; Menschaert and Fenyo 2015). The need for computational solutions to apply proteogenomics more broadly has been noted (Castellana and Bafna 2010; Renuse et al. 2011; Armengaud et al. 2014; Nesvizhskii 2014). Of particular interest are tools that create customized databases (DBs) to identify evidence for unannotated ORFs. RNA-seq data have been used to limit the protein search DB size to achieve better statistical power (Wang et al. 2012; Woo et al. 2013; Zickmann and Renard 2015). Other MS-friendly DB solutions that integrate data from different species or strains include MScDB (Marx et al. 2013), MSMSpdbb (de Souza et al. 2010), and PG Nexus (Pang et al. 2014). Even pipeline solutions were developed that allow to search proteomics data against a six-frame translation based DB, including Peppy (Risk et al. 2013), Genosuite (Kumar et al. 2013) and PGP (Tovchigrechko et al. 2014). However, an integration that leverages benefits of manually curated reference annotations and a six-frame translation into one highly informative, non-redundant and transparent resource has not been accomplished so far.

Here, we address this unmet need of the microbiology and proteomics community and present a strategy that takes the MS-friendly DB concept one important step further. Our solution integrates and consolidates CDSs from multiple reference annotation resources, *ab initio* gene prediction algorithms and *in silico* ORFs (a six-frame translation, also considering alternative start codons). Identifiers capturing information about overlap and differences among the resources are created, as well as a GFF (generic feature format) file storing all annotations and a highly informative, integrated proteogenomics search database (iPtgxDB). Based on an extension of PeptideClassifier’s concept of unambiguous peptides (Qeli and Ahrens 2010) for prokaryotes, close to 95% of the peptides in this search DB unambiguously identify one distinct protein sequence. This greatly facilitates downstream analysis by overcoming the need to dis-entangle protein groups implied by shared peptides. As a first proof-of-concept, searching data from a complete, condition-specific expressed proteome against a *Bartonella henselae* iPtgxDB allowed to uncover novel ORFs uniquely predicted by each of the resources, illustrating the value of this integrated approach. Of note, the expression of the large majority of novel ORFs could be confirmed by independent targeted parallel reaction monitoring (PRM) MS. Our approach is flexible: iPtgxDBs can be created both for model organisms with readily available reference annotations and for newly sequenced genomes. We illustrate this benefit using a completely assembled genome of a laboratory strain and proteomics data to track evidence for unique, differentially expressed proteins down to single amino acid variations (SAAVs). iPtgxDBs were also generated and evaluated for *B. diazoefficiens* and *E. coli,* and the software to create such DBs for any prokaryote is released. As open source platform (https://iptgxdb.expasy.org), iPtgxDBs enable many research groups to take full advantage of completely sequenced genomes by improving genome annotations with proteogenomics.

## Results

### Experimental evidence underscores the need for a general proteogenomics approach

We used the α-proteobacterium *Bartonella henselae* strain Houston-1 (Bhen) to explore how genome annotation differences could best be integrated for a proteogenomics approach. A comparison of four Bhen reference genome annotations and results from two *ab initio* gene prediction tools (see Methods) confirmed reports for other organisms (Poole et al. 2005; Bakke et al. 2009; Cuklina et al. 2016) that both the number of predicted ORFs and their precise start sites largely differ (see Figure S1). Only 50% of the Bhen CDSs were annotated or predicted completely identical by all six resources and 37% were unique to one resource (Figure S1A). Of note, 23% of the CDSs of the recent NCBI RefSeq2015 re-annotation differed from RefSeq2013 (Table S1): 55 CDSs were removed (99 added), 74 CDSs were shortened (54 extended), and 64 pseudogenes were removed (15 added).

To assess the validity of the RefSeq2015 re-annotation, we relied on an ideal dataset: a complete prokaryotic proteome (including many low abundant proteins) expressed under two conditions that mimic those encountered by Bhen in the arthropod vector midgut (uninduced condition) and the bloodstream of its mammalian host (induced condition), which had been searched against RefSeq2013 (Omasits et al. 2013). A search against a RefSeq2015 protein DB provided experimental evidence for many of the re-annotations, including 6 sORFs that we had previously identified with a prototype of our proteogenomics approach as novel (Table S1), and which have since been added to RefSeq2015. Also, among the 55 removed CDSs, we found 32 of 52 proteins we had earlier singled out as potential over-predictions (Omasits et al. 2013). However, we also found several cases that supported the earlier RefSeq2013 annotation, including expression evidence for CDSs that were relabeled as pseudogenes and for removed CDSs (Table S1). This highlights the need for an integrated, yet general approach to address this fundamental problem of gene annotation inconsistency.

### A general, integrative proteogenomics approach

An ideal solution to capture the full protein-coding potential of genome sequences should therefore i) consider results from different reference genome annotations (Nesvizhskii 2014), which often include substantial manual curation efforts from experts, and from *ab initio* gene prediction tools, ii) allow to identify the small fraction of true functional sORFs often missed by the above annotations or predictions, iii) aid in the annotation of newly sequenced genomes, and iv) enable scientists to visualize their experimental proteomics results in the context of both the genome and all available annotations.

To our knowledge, existing tools only address a subset of these requirements. These include pipeline solutions that rely on a six-frame translated genome like Peppy (Risk et al. 2013), which aims to improve the scoring function for peptide spectrum matches (PSMs), Genosuite (Kumar et al. 2013), which uses four distinct search algorithms before integrating and visualizing the results, and PGP (Tovchigrechko et al. 2014), which draws on the experience of many proteogenomics studies (Venter et al. 2011) and highlighted the need for stringent criteria to accept novel ORFs. However, these tools do not integrate different annotation sources. Some MS-friendly integrated DBs accomplish this, such as MScDB (Marx et al. 2013), which uses a peptide-centric clustering algorithm to combine e.g. cross-species DBs, or MSMSpdbb, which allows to create a non-redundant protein DB for multiple closely related bacterial strains (de Souza et al. 2010). However, they do not integrate different annotations of the same genome. PG Nexus (Pang et al. 2014) uses the NCBI RefSeq annotation, a Glimmer *ab initio* prediction (Delcher et al. 2007) and a six-frame translation against which peptides are searched with Mascot and later visualized onto the genome. However, the annotations are not integrated and consolidated; the boundaries of novel ORFs still have to be discovered based on peptide evidence, which requires substantial manual effort. In addition, Mascot is not ideal for the task of identifying novel ORFs in proteogenomics approaches (Omasits et al. 2013; Risk et al. 2013). To address all of the above objectives in one integrated solution, we devised a proteogenomics workflow that relies on three steps (Figure 1).

**Figure 1.**
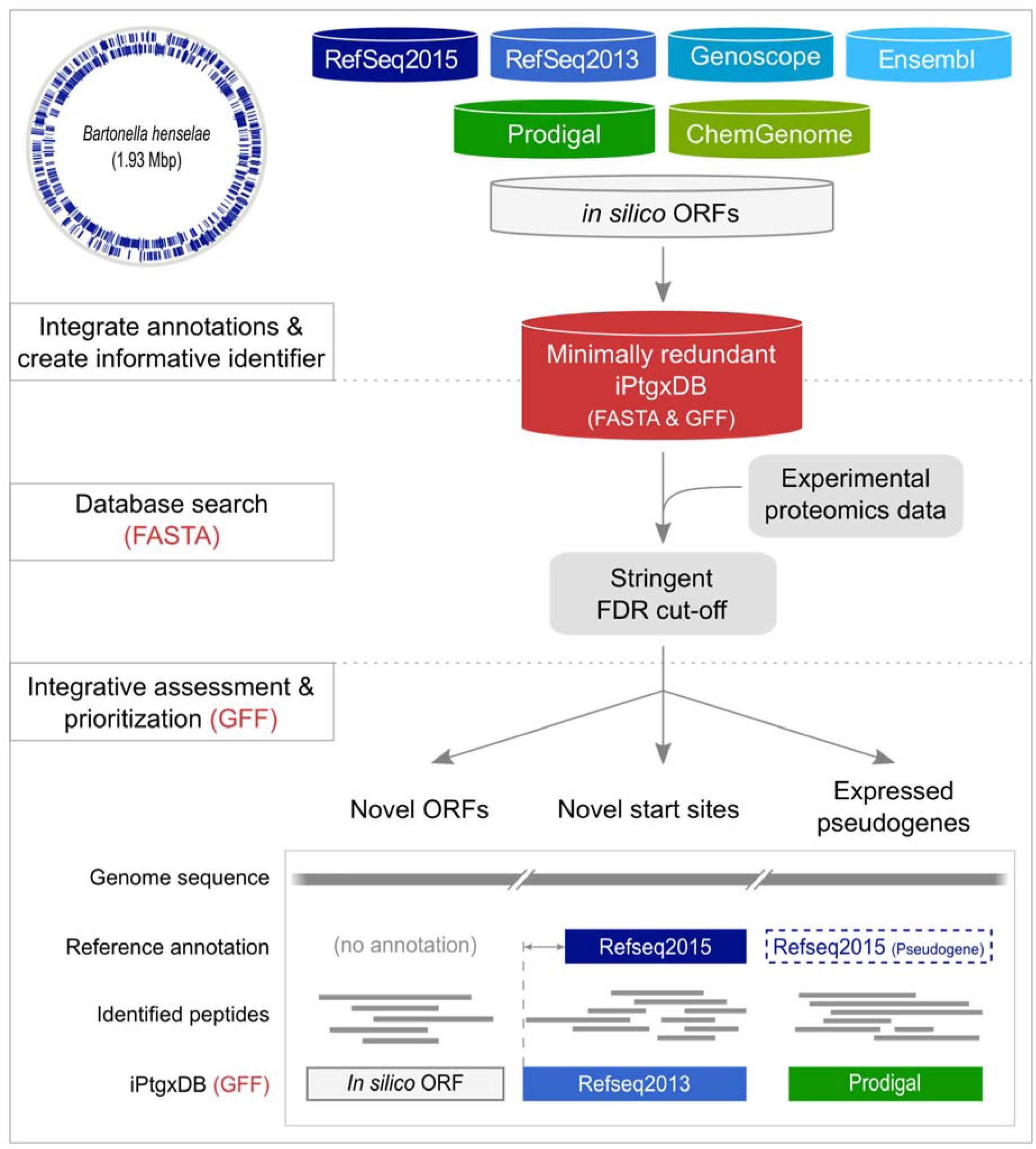
Integrative proteogenomics workflow. For a completely sequenced prokaryotic genome (Bhen is shown as example with annotated CDSs), reference genome annotations (blue containers), results from *ab initio* gene prediction algorithms (green containers), and *in silico* ORFs (white container), are downloaded or computed and integrated in a first pre-processing step (upper panel). All CDS and pseudogene annotations are matched, informative gene identifiers are created and stored in a minimally redundant iPtgxDB (red container; searchable protein sequences in FASTA format, integrated annotations in GFF format). Experimental proteomics data are matched to the DB using a target-decoy approach relying on stringent FDR cut-offs (middle panel). Identified PSMs and peptides are post-processed to visualize novel candidates (lower panel) in the context of experimental data integrated with the GFF file.

First, popular reference genome annotations, the results of *ab initio* prediction algorithms, and *in silico* ORFs from a six-frame genome translation were combined into an integrated proteogenomics search database (iPtgxDB) (Figure 1, upper panel) with the aim to capture the entire genomic protein-coding potential (see Methods). On top of genome annotations from NCBI RefSeq, Ensembl and Genoscope, we included results of the *ab initio* gene prediction algorithms Prodigal (Hyatt et al. 2010), which performs well even for genomes with high GC content where gene calling is more difficult (Marcellin et al. 2013), and ChemGenome. The latter relies on physico-chemical characteristics of codons calculated by molecular dynamics simulations (Singhal et al. 2008) and is thus quite different from Prodigal and similar tools (Pati et al. 2010). Finally, to be able to identify functional sORFs, which are often missed due to rather conservative length thresholds for *ab initio* predicted ORFs, all potential *in silico* ORFs (a modified six-frame translation; see Suppl. Methods) above a selectable length threshold were added. A literature search for experimentally validated prokaryotic sORFs (Zuber 2001; Rowland et al. 2004; Venter et al. 2011) revealed that novel sORFs were longer than 20 aa, with very few exceptions (Hemm et al. 2008). To balance comprehensiveness and avoid loss of statistical power when searching large DBs (Blakeley et al. 2012; Noble 2015), we selected a length threshold of 18 aa.

In a second step, proteomics data - ideally comprehensive expression data obtained under multiple conditions (Ahrens et al. 2010) - is searched against the iPtgxDB and stringently filtered (Figure 1, middle panel). We used the search engine MS-GF+, which rigorously computes E-values of PSMs based on the score distribution of all peptides (Kim and Pevzner 2014) and which had performed favorably in our hands for large shotgun proteomics datasets (Omasits et al. 2013) as well as in proteogenomics studies (Risk et al. 2013; Zickmann and Renard 2015; Cuklina et al. 2016).

In a third step, peptide evidence is visualized in the context of the genome and all annotations (contained in a GFF file) using a genome browser such as IGV (Robinson et al. 2011) (Figure 1, lower panel). Candidates in major classes of novelty include novel ORFs, different or additional start sites, and expressed pseudogenes. These can be inspected in the context of experimental data (e.g. proteomics and transcriptomics data), functional annotations and other features to enable a comprehensive assessment and prioritization.

### Creating minimally redundant but maximally informative protein search databases

A unique aspect of our proteogenomics approach is that almost all MS-identifiable peptides of the iPtgxDB unambiguously identify one specific protein (Figure 2). To achieve this, we extended our PeptideClassifier concept (Qeli and Ahrens 2010) for prokaryotes. PeptideClassifier was developed to classify the information content of peptides with respect to their originating gene model(s) into six classes; class 1a peptides are most informative and allow unambiguous identification at the protein sequence, protein isoform and gene model level (Figure S2). Our extension for prokaryotes now treats protein sequences with a common stop codon and varying start positions (N-termini) as a protein annotation cluster, i.e. variants of a prokaryotic gene model (similar to isoforms of a eukaryotic gene model). Class 1a peptides remain most informative as they are unique to one entry in a DB, while class 1b peptides map uniquely to one annotation cluster with all identical sequences. Class 2a peptides identify a subset of sequences from an annotation cluster and class 2b peptides map to all sequences of an annotation cluster. Class 3a peptides map to identical sequences from different annotation clusters (typically duplicated genes). Class 3b peptides map to different sequences from different annotation clusters and are least informative.

**Figure 2.**
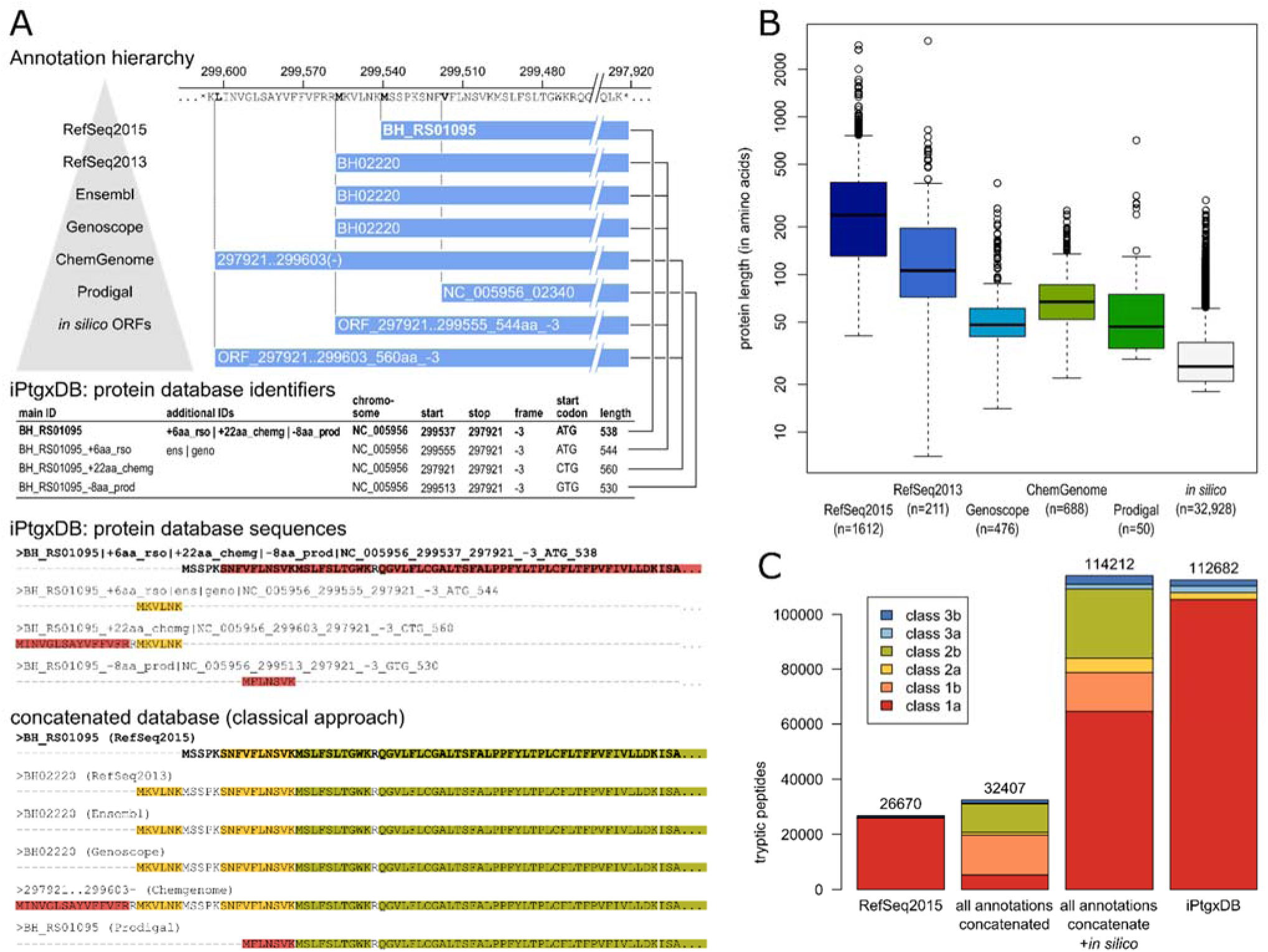
Generating an iPtgxDB with informative identifiers and a minimally redundant protein search DB in FASTA format. (A) CDSs and pseudogenes of 7 resources are integrated in a stepwise fashion. Informative protein identifiers are created, illustrated for the annotation cluster with the RefSeq2015 anchor sequence BH_RS01095 shown in bold, where three additional start sites exist. The four different proteoforms are added to the protein search DB: the anchor sequence (bold) with the full protein sequence, the extensions (RefSeq2013 and ChemGenome) add the upstream sequence up to the first tryptic cleavage site within the anchor sequence. The shorter Prodigal prediction uses an alternative start codon resulting in a distinguishable N-terminal peptide, and therefore gets also added. The two *in silico* ORFs are identical to annotations higher up in the annotation hierarchy, and therefore are not added. Peptide classes are shown for the N-terminal sequences of the CDS annotation cluster (see also Fig. 2C). (B) Boxplots of protein length for RefSeq2015 and of those proteins that get added in each successive step to the protein search DB illustrate that we include many sORFs potentially missed in the reference annotations. (C) Bar chart showing the DB complexity and the peptide classes for RefSeq2015, all 6 integrated annotations without and with *in silico* ORFs, and the final iPtgxDB. The legend shows colors for the six peptide classes.

The stepwise integration of resources, carried out in Figure 2A for Bhen as a model, follows a hierarchy: to leverage the quality of manual curation efforts we start with reference annotations, then *ab initio* predictions, then *in silico* ORFs. The anchor sequence is selected from the annotation highest up in the hierarchy, i.e. here RefSeq2015, unless no CDS is predicted in a given genomic region. Each subsequent resource added new protein clusters, and new extensions or reductions (i.e. alternative start sites) to an existing cluster, while identical annotations are collapsed (Table 1). Alternative start codons are captured with our approach, even for *in silico* ORFs (Figure 2A).

**Table 1.**
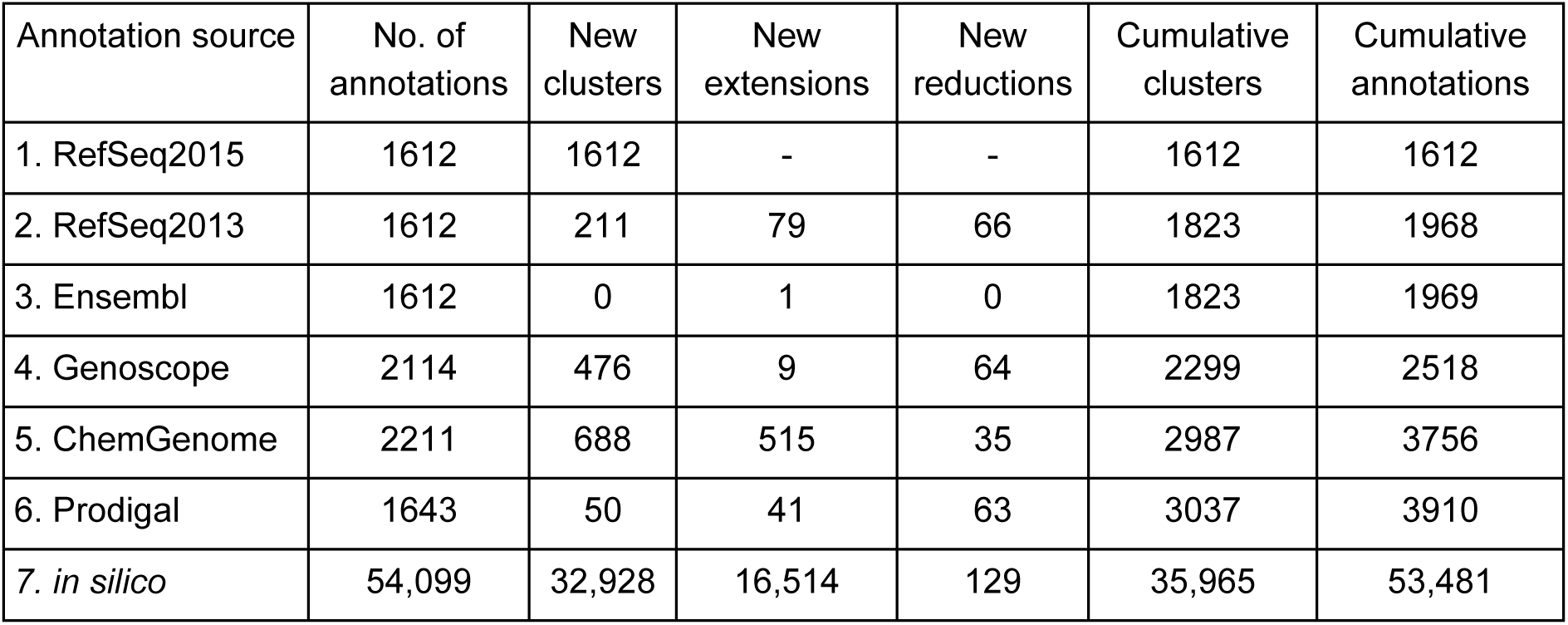
Result of the stepwise, hierarchical integration of resources for Bhen. The number of annotations per source, new protein clusters, new extensions and reductions are shown for each step and summarized under cumulative clusters and cumulative annotations. Overall, protein sequences for 53,481 annotations mapped to 35,965 annotation clusters. The final iPtgxDB had 51,541 entries, as sequences < 6 aa (1750), i.e. not identifiable with shotgun proteomics, and indistinguishable internal start sites (190) were not considered.

The four distinct sequences (collapsed from eight annotations) of the protein cluster for apolipoprotein N-acyltransferase with the anchor sequence BH_RS01095 (Figure 2A) illustrate this key principle: by only adding the full sequence of the anchor protein plus class 1a peptides (red in Figure 2A) that can unambiguously identify extensions or internal start sites and few class 2a peptides (yellow in Figure 2A) to the iPtgxDB, we minimize redundancy and maximize information content of the peptides, compared to adding all protein sequences (Figure 2A, lower panels). In the latter case, many peptides classified as 2a or 2b, which imply a subset or all annotations of a CDS cluster, would get added to the iPtgxDB. Identification of such shared peptides greatly impedes downstream analysis.

The protein identifiers of the four distinct sequences transparently capture overlap and differences of the annotations (see Methods); they show in which resource(s) the identified CDS is annotated, if and how the annotations differ, whether it is a novel ORF or an alternative start site, again largely improving downstream data analysis (Figure 2A).

The identifiers also contain genomic coordinates, allowing to visualize all experimental peptide evidence for a novel ORF in its genomic context along with all integrated annotations provided in the iPtgxDB GFF file. Peptides implying any other sequence (e.g. one of the three identifiers below the anchor sequence identifier in Figure 2A) would inform the experimentalist at a glance that novel information compared to RefSeq2015 was uncovered (see Supplemental Methods for examples how to “interpret” the identifiers).

A box plot of the lengths of the proteins added to our DB in the stepwise processing illustrates that we capture increasingly smaller proteins. Adding *in silico* ORFs down to a selectable length threshold allows us to query the entire protein-coding potential of the genome (Figure 2B).

The final Bhen iPtgxDb contains 51,541 entries (Table 1). Importantly, 94% of all theoretically MS-identifiable tryptic peptides (6-40 aa) allow unambiguous identification of one protein, i.e. class 1a peptides (red, Figure 2C). For RefSeq2015, almost all peptides mapped uniquely to one protein, which is common for a prokaryote (Figure 2C). Combining all 6 annotations resulted in a modest increase of tryptic peptides (23%); however, most peptides (85%) now matched at least two annotations: either annotations for an identical sequence (class 1b) or annotations of proteins with different length, but of the same annotation cluster (class 2a or 2b, i.e. a subset or all of the proteoforms of the cluster), which would greatly complicate the interpretation of proteomics search results. Adding *in silico* ORFs significantly increased the number of peptides, adding mainly new unique peptides (class 1a) for ORFs in regions without annotation (Figure 2C). Our careful integration collapses identical sequences and removes 1b and 2b peptides from the iPtgxDB.

Defining DB complexity as number of distinct tryptic peptides of 6-40aa length, the complexity of the resulting iPtgxDB was approximately 50% of that of a full six-frame translated genome that Mascot (Perkins et al. 1999) or PG Nexus (Pang et al. 2014) would rely on to identify proteogenomic evidence for novel peptides. Despite the relatively large number of entries, the DB complexity is only 70% of that of baker’s yeast and below 20% of a human protein DB (Table S2).

### Searching Bhen proteomics against our iPtgxDB identifies novel ORFs

We next searched existing data from a comprehensive expressed proteome, an *in vitro* model mimicking interaction of Bhen with the arthropod vector (uninduced condition) or its mammalian host (induced condition) (Omasits et al. 2013), against the Bhen iPtgxDB using MS-GF+. Relying exclusively on unambiguous class 1a peptides, this allowed to systematically identify expression evidence for novel ORFs, novel start sites, and CDSs wrongly annotated as pseudogenes (Table 2). Importantly, each of the reference genome annotations, *ab initio* gene prediction tools, and *in silico* predicted ORFs provided unique novel hits, underlining the value of our integrated approach (Table 2, Figure S3). These hits are novel compared to the most common approach of using the latest reference annotation as search DB, i.e. RefSeq2015 in this case.

**Table 2.** Summary of novel information uncovered by the integrated proteogenomics approach. Compared to RefSeq2015, each resource added some novelty with respect to Bhen’s overall protein-coding potential. Overall, close to 80% of the identified novelties could be independently confirmed by PRM (Table S3).

| Annotation source | Novel protein-coding ORF |  | Evidence for alternative NH2-terminus |
| --- | --- | --- | --- |
|  | not in RefSeq2015 | pseudogene in RefSeq2015 |  |
| RefSeq2013 / Ensembl | 4 | 10 | 5 |
| Genoscope | 4 | 1 | 0 |
| ChemGenome | 1 | 2 | 5 |
| Prodigal | 3 | 1 | 2 |
| *in silico* ORFs | 10 | 1 | 5 |
| total | 22 | 15 | 17 |

When searching large datasets, it is imperative to use stringent cut-offs. This is particularly relevant for proteogenomics, where correctly identified novel information would require a genome annotation change. We relied on an estimated PSM-level false discovery rate (FDR) cut-off of 0.01%, which resulted in a peptide-level FDR of 0.12%. This cut-off is about 10-fold more stringent than in other proteogenomics studies (Krug et al. 2013; Kumar et al. 2013; Chapman and Bellgard 2014; Zickmann and Renard 2015), and closer to the cut-offs used by Payne and colleagues (peptide level FDR cut-off 0.3%) (Venter et al. 2011). Of particular note, the E-value score distribution of PSMs that identify novel features is also bi-modal, similar to that of PSMs identifying annotated proteins in the target DB (Figure 3A). To claim a potential novel ORF, we required at least 3 PSMs to class 1a peptides if predicted by a reference genome annotation/*ab initi*o prediction tool, and 4 PSMs to class 1a peptides for *in silico* ORFs, in line with earlier recommendations (Nesvizhskii 2014). Furthermore, all genomic regions encoding novel candidates/start sites were expressed (Table S3).

**Figure 3.**
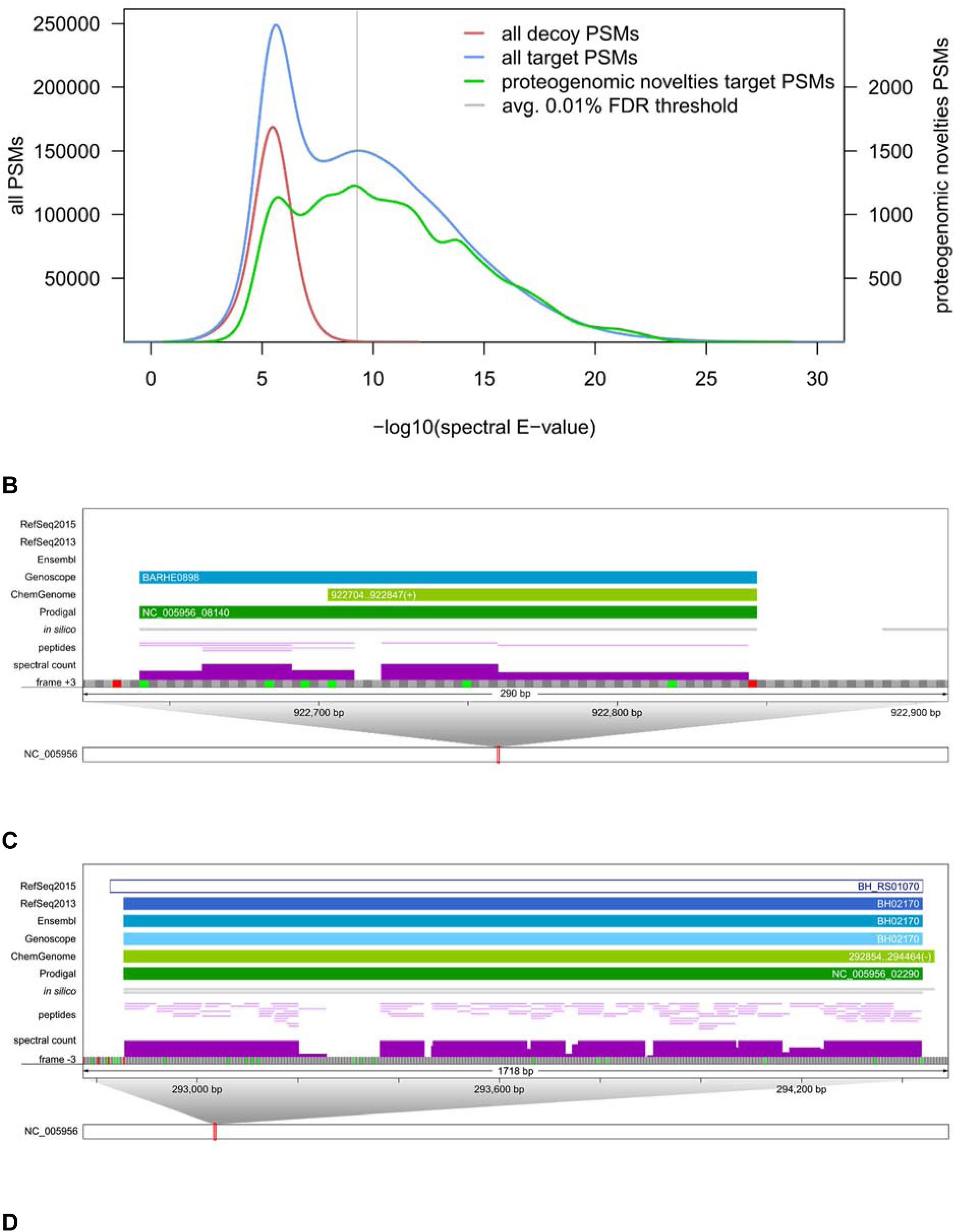
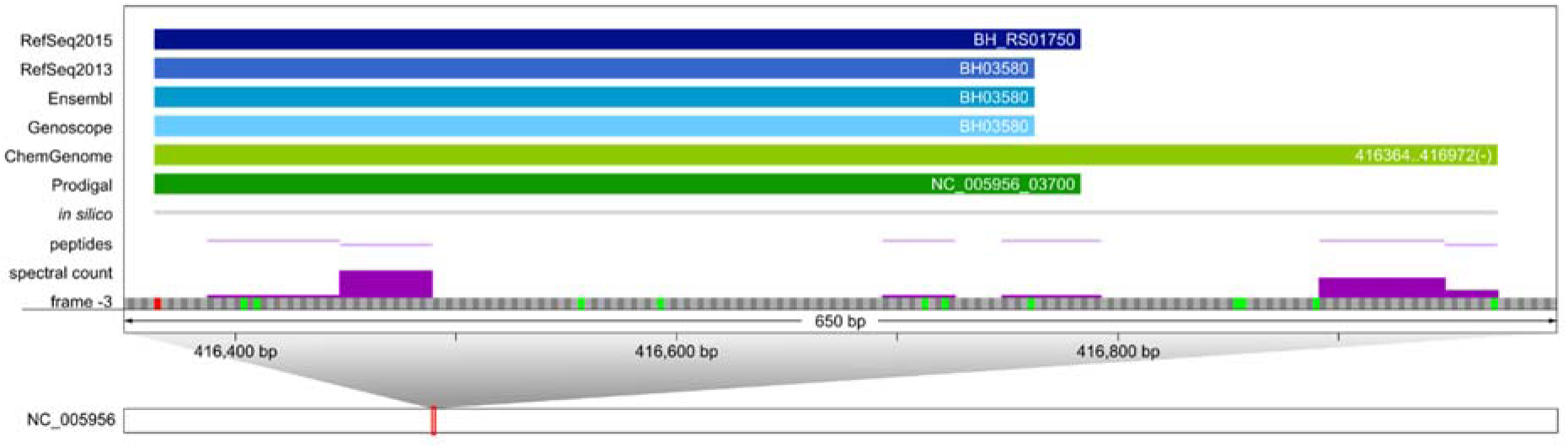
Examples of novel information uncovered by integrative proteogenomics. (A) E-value distribution of PSMs against proteins of decoy and target DB (red and blue lines, left scale) plus the distribution of those PSMs that uncovered novelties (green line, right scale). A PSM level FDR cut-off of 0.01% was selected per sample. (B-D) Zoomed in views of genomic regions that harbor novelties. For illustration, a single frame of the forward/reverse strand with possible start (green) and stop codons (red) is shown, along with annotations and experimental evidence (spectral counts scaled from 0 to 20). (B) Example of a novel sORF of 68 amino acids (BARHE0898, frame +3). (C) Example of a highly expressed pseudogene (RefSeq2015: BH_RS01070, frame -3). 2244 spectra are mapped to 117 peptides of NusA, which is annotated as pseudogene in RefSeq2015 for unknown reason. There is no experimental evidence for the +8 aa N-terminal extension predicted by ChemGenome. (D) Proteomic expression evidence supports a 63 aa longer proteoform of BH_RS01750 (frame -3) uniquely predicted by ChemGenome.

Overall, 37 novel Bhen ORFs (with respect to RefSeq2015) were identified (Table 2, Figure S3): 12 annotated by another resource or *ab initio* prediction tool, 10 *in silico* only predicted ORFs, and 15 with a pseudogene annotation. In addition, 17 alternative start sites were identified. The median length of 22 novel ORFs (excluding pseudogenes) was 48 aa, that of 6 novel ORFs previously identified with a prototype versus RefSeq2013 (i.e., prior to the re-annotation; Table S1) was 80 aa (Figure S4). This confirms that the novel ORFs represent sORFs commonly underrepresented in genome annotations. Analysis of the estimated expression levels of the novel ORF candidates including pseudogenes (see Methods) furthermore indicated that several of the sORFs are well expressed proteins (Figure S4) that may carry out important functions.

Examples of novel ORFs included differentially expressed sORFs such as BARHE0898 (68 aa), that was expressed roughly 6.5-fold higher in the induced condition (Table S3). Peptide evidence supported the longer form of this ORF annotated by Genoscope and Prodigal (Figure 3B). Another novel sORF of 67 aa, a lipoprotein uniquely predicted by ChemGenome, was only identified in the uninduced condition (Figure S5A). Even smaller ORFs were identified, including a well-expressed (9 peptides, 143 PSMs) sORF of 49 aa that was uniquely predicted by Prodigal (Fig S5B, Table S3), and an *in silico* only predicted sORF of 34 aa (Figure S5D). We also identified a highly expressed RefSeq2015 pseudogene (BH_RS01070, frame -3, Figure 3C) annotated as normal CDS (transcription elongation factor NusA) by RefSeq2013, Ensembl, and Genoscope. Other mis-annotated pseudogenes included a potassium-efflux transporter (BH10840) and the *Bartonella* effector protein BepD (BH13410, Table S3). Finally, for BH_RS01750 a hypothetical protein encoded in a prophage region, only ChemGenome correctly predicted a 63 aa longer proteoform than annotated by other resources; its expression was supported by several peptides (Figure 3D). Proteomic data thus can support novel start sites (even multiple start sites; Figure S5C) and distinguish between those predicted by different reference annotations.

### Confirmation of novelties by independent targeted proteomics

In order to confirm novel ORFs by independent methods, the expression of novel candidates at the protein level was assessed by targeted proteomics experiments (Figures 3 B-D and S5 A-D) with parallel reaction monitoring (PRM) assays (Peterson et al. 2014). For this highly sensitive method, cytoplasmic (cyt) and total membrane (TM) extracts were prepared from new biological samples as described (Omasits et al. 2013) (see Methods). Overall, we were able to validate 107 of 138 targeted peptides (78%; Tables S3, S4), including low expressed novel proteins implied by 1 peptide and 3 PSMs. We had previously derived predominant subcellular localizations (SCL) for all proteins (Stekhoven et al. 2014), which we computed here also for the novel candidates (see Supplemental Methods, Table S3). Importantly, the SCL data agreed with the PRM evidence in either cyt or TM fractions, thereby adding yet another layer of support to the confirmed novel ORFs (Table S3). The validation success was 100% for the 6 novel ORFs identified previously by our prototype (novel with respect to the RefSeq2013 annotation, Figure S4), around 80% for 15 expressed pseudogenes and 12 novel ORFs from another genome annotation/prediction, 60% for novel *in silico* ORFs, and around 55% for novel start sites (TableS3).

Of note, we overall identified 38 of 51 lipoproteins predicted to have a SpII cleavage site (LipoP, version 1.0) (Table S5). Two of these were identified among the 12 novel ORFs (one predicted by Genoscope, one by ChemGenome), and two others among the 6 novel ORFs identified previously, which have since been incorporated in Refseq2015. All four candidates were validated by PRM (Table S3), and their predominant SCL indicated that they were found exclusively in the total membrane or outer membrane fractions (Table S3). Lipoproteins could thus represent a class of proteins for which a substantial percentage is missed in reference genome annotations, which is relevant given their important roles in signaling, protein folding and export, virulence, immunity and antibiotic resistance (Kovacs-Simon et al. 2011).

### *De novo* assembly and genome comparison of Bhen strains underlines the importance of a correct genome sequence

Massively reduced sequencing costs and improved assembly algorithms make it possible to determine the actual genome sequence of key bacterial strains used in a laboratory, which provides an optimal basis to integrate functional genomics data and to correctly identify novel sORFs. We thus explored to what extent our lab strain (MQB277) differed from the Bhen reference strain (Alsmark et al. 2004), and whether we could detect protein expression evidence for novel ORFs in unique genomic regions, and for SAAVs. Conceptually, this allowed to test our approach on a newly sequenced genome, now relying on an iPtgxDB integrating Prodigal predictions plus *in silico* ORFs, but without curated reference genome annotations.

We *de novo* assembled (Chin et al. 2013) the PacBio sequenced genome of the MQB277 lab strain, derived from Bhen CHDE101, a Bhen variant-1 strain (Lu et al. 2013) into one 1,954,773 bp high quality contig (see Suppl. Methods), i.e. ~23.7 kbp longer than the NCBI reference genome (Figure 4A). To compare closely related genomes in the context of experimental evidence, we devised a “virtual genome” concept, i.e. a coordinate system that integrates sequences from reference genome and *de novo* assembly (Figure 4). This allowed us to integrate annotation and experimental data tracks, to efficiently zoom down to the single nucleotide level and to inspect all lines of evidence for observed differences.

**Figure 4.**
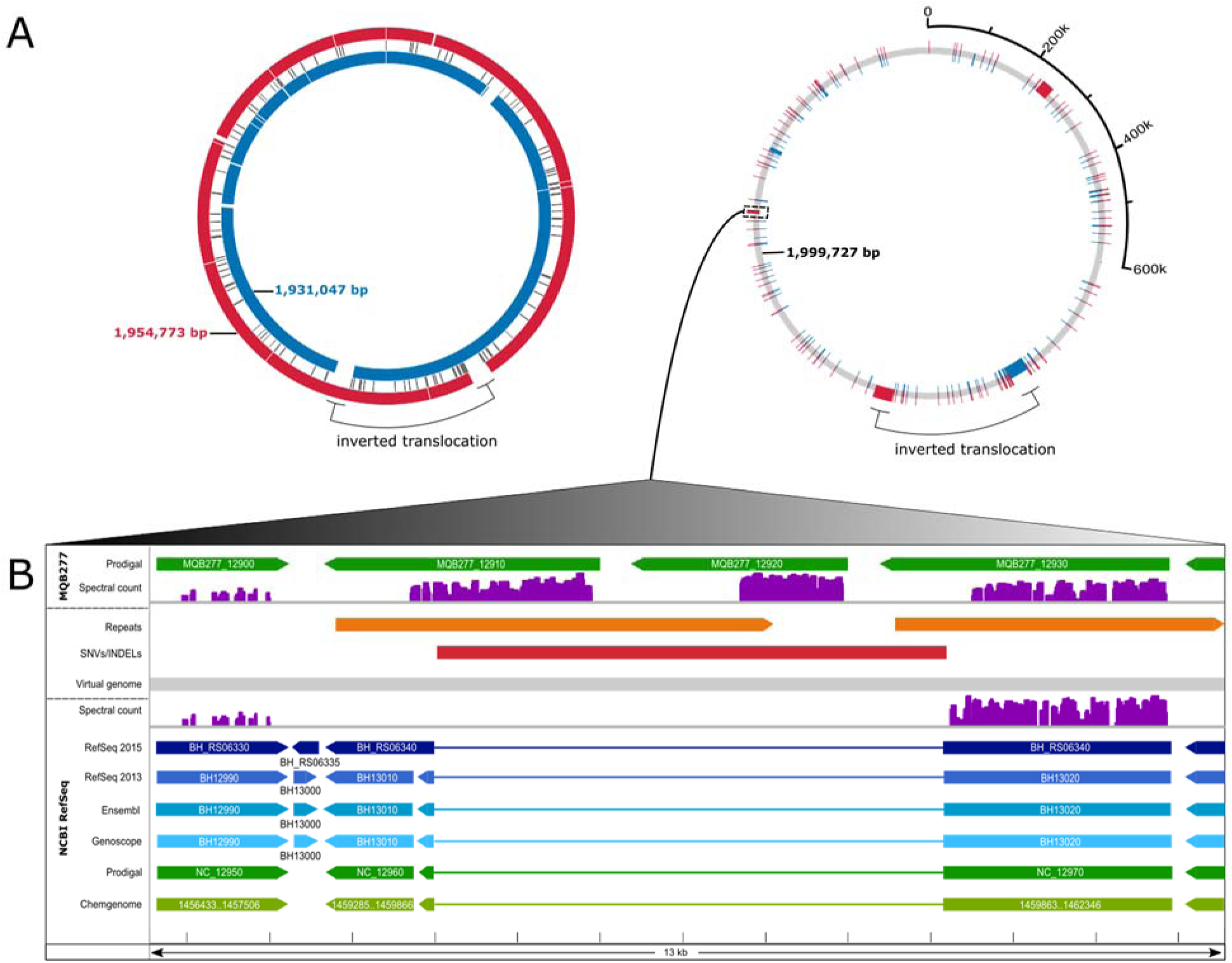
Comparative analysis in the context of experimental data. Integrated visualization of two closely related genomes through a virtual genome concept. (A) On the left, the Bhen NCBI RefSeq genome (inner blue circle) is aligned to our *de novo* assembly (outer red circle). A large inversion-translocation (black bracket) is marked; several insertions or deletions in either genome are shown (white spaces) and a center track for single nucleotide variations (SNVs). To the right, the virtual genome (gray) is shown which incorporates both genome sequences including all differences into a common coordinate system. Unique sequences are shown in blue or red, respectively (the inversion-translocation present in both genomes is left as is). (B) Zoom into the region harboring a 6088 bp insertion in MQB277 (red bar), showing annotations for RefSeq genome (below the virtual genome track) and assembly (above the virtual genome track), plus experimental proteomics evidence mapped against both genomes (spectral count scaled from 0-800). This region harbors a direct repeat only in the assembly (orange bars). Three CDSs (MQB277_12910, MQB277_12920, MQB277_12930) annotated as autotransporters are highly expressed; the first two (novel CDSs) are only detected (unambiguous 1a peptides) with the correct genome sequence available.

Overall, we noted a large inversion translocation (34.4 kbp) close to the terminus of replication, previously reported for some *Bartonella* isolates (Lindroos et al. 2006), and three insertions of 22.1, 6.1 and 1.4 kbp in the MQB277 assembly (Table S6). The 22.1 and 1.4 kb insertions affected a genomic region encoding the surface protein BH01510, a BadA1 adhesin (Figure S6), and major pathogenicity factor that mediates binding of *B. henselae* to extracellular matrix proteins and endothelial cells (Riess et al. 2004). A complex repeat structure in this region of the assembly harbored additional ORFs, whose expression was supported by unambiguous peptide evidence. In line with the CDS MQB277_01630 lacking a C-terminal membrane anchor, its predominant SCL was cytoplasmic (for more detail, see Figure S6). These data help to explain earlier experimental data that had demonstrated lack of or much lower BadA surface expression in the Bhen CHDE101 variant strain (Lu et al. 2013).

Of note, the 6.1 kbp insertion harbored two CDSs predicted to encode autotransporter proteins (Figure 4B). Both MQB277_12910 and MQB277_12920 were unique to the high quality PacBio assembly, which contains a direct repeat in this region (missing in the reference), indicative of a duplication event. Searching against the MQB277-based iPtgxDB, all 3 proteins were highly expressed (Figure 4B, MQB277 track), compared to only BH_RS06340 based on the RefSeq2015 protein DB (NCBI RefSeq track). Importantly, the novel CDS MQB277_12910 was among the most upregulated (42-fold) proteins in the induced condition (Table S3), even higher than the autotransporter BH_RS06340 (38-fold). This data is in line with the dramatic re-organization of the membrane proteome reported earlier (Omasits et al. 2013). Our SCL data indicated that all three proteins were localized in the outer membrane (Table S3).

The assembly also comprises a genomic region harboring a 1 bp insertion and a 81 bp deletion with respect to the reference. Because of the frameshift caused by the insertion, protein expression of a CDS annotated as ABC transporter (downstream of the insertion) was only observed in the lab strain assembly (Figure 5A, Table S6), which is also supported by transcriptomics data (see Methods). Due to the frameshift, CDSs in this region were either annotated as pseudogenes or as split CDSs (NCBI RefSeq track). This example is furthermore noteworthy, as our transcriptomics dataset (Omasits et al. 2013) had been re-analyzed with an RNA-seq based proteogenomics approach (Zickmann and Renard 2015).

**Figure 5.**
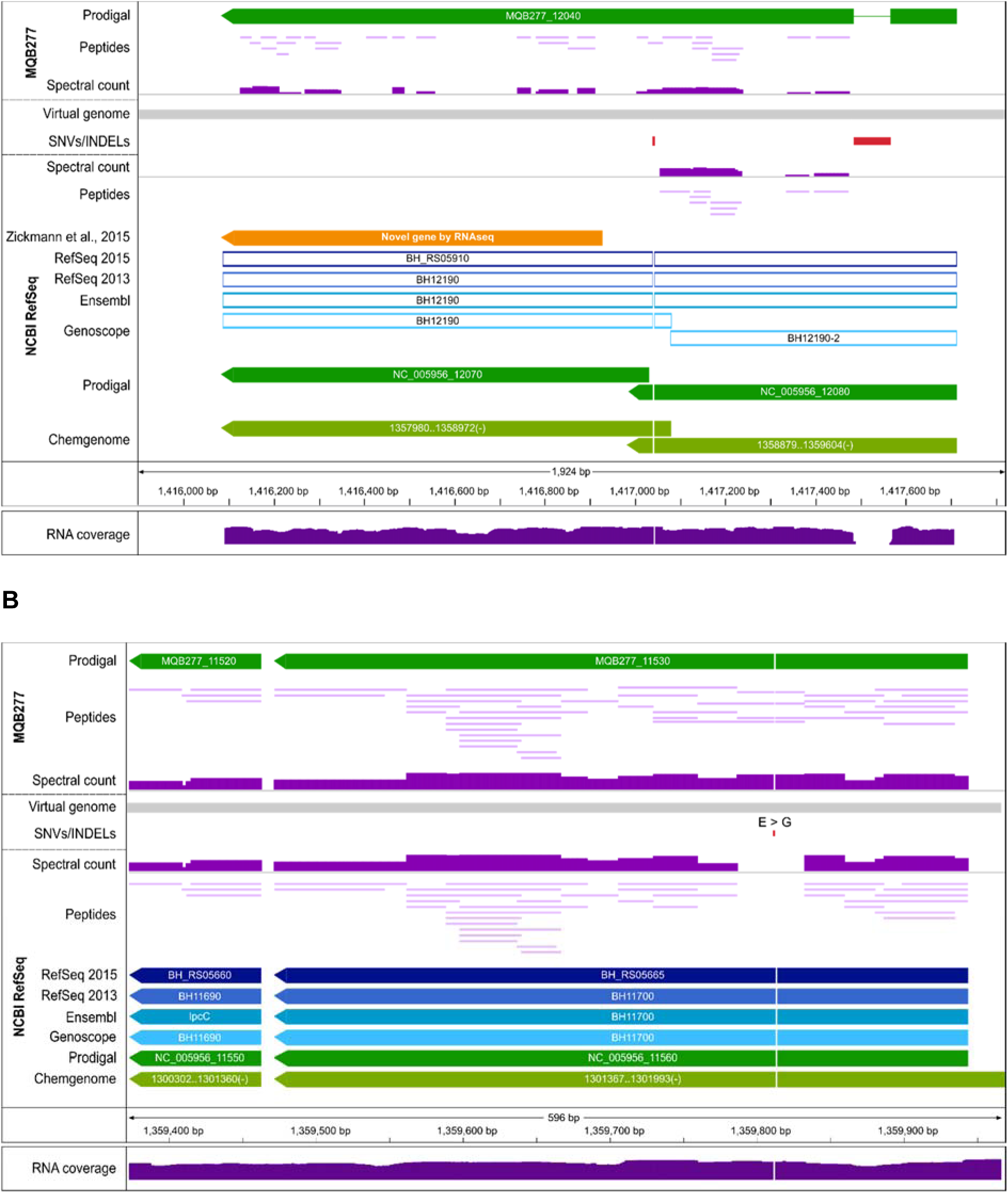
Protein evidence for SAAVs. (A) Genomic region encoding an ABC transporter (BH_RS05910). RefSeq and Ensembl annotate it as pseudogene, Genoscope as fragmented pseudogene, while Prodigal and Chemgenome predict 2 CDSs. The reference genome (below gray virtual genome bar; NCBI RefSeq track) differs from the MQB277 assembly (MQB277 track above the virtual genome) by an insertion of 81 bp and a 1 bp deletion (red boxes); the 1 bp deletion causes a frameshift, evidenced by the lack of protein expression downstream of it (spectral count below the virtual genome; scaled from 0 to 800) and by transcriptomic data (reads mapped to the reference genome all support the insertion; lower panel). In contrast, the protein encoded by MQB277_12040 in the assembly is expressed over almost its entire length (class 1a peptides; 1 peptide identified by 7 PSMs spans the frameshift region), also supported by transcriptomic reads mapping without any mismatch (data not shown). (B) Evidence for a SNV causing a non-synonymous SAAV in the CDS of transcription elongation factor GreA. Four peptides (2, 4, 8, 39 PSMs) confirming this SAAV (Glycine in reference to Glutamic acid in our assembly) are mapped to this position in MQB277.

Two novel ORFs were reported, one of which is shown in Figure 5A (orange arrow). However, only with the correct assembly at hand can novel ORFs be identified completely accurately. Our integrated analysis demonstrates that the novel ORF in question is in fact longer, and its expression is supported by class 1a peptides (MQB277 track). Of note, we found several examples where SNVs led to non-synonymous protein sequence differences, as e.g. for transcription elongation factor GreA (Figure 5B). The expression of this and 11 additional SAAVs (Table S3) was again independently validated by PRM assays. Analysis of the 274 SNVs observed between the two genomes indicated that they were significantly enriched in a limited number of regions encoding surface proteins (114/274), including 4 of 8 hemagglutinins, and 4 hemolysin activator proteins (HECs) (Table S6). Together, our data provide multiple lines of evidence for the earlier postulation that genome rearrangements observed in natural Bhen populations affect variation of surface proteins (Lindroos et al. 2006).

Finally, the search against the assembly-based protein DB led to overall 10,410 more assigned PSMs, and 441 peptides (same 0.01% PSM level FDR threshold; Table S7). Overall, these results emphasize the value that research groups can gain by sequencing and assembling their most important strains.

### Integrated proteogenomics approach is generically applicable

Genome annotation is more difficult and error-prone for genomes with high GC content; they contain more spurious ORFs and fewer stop codons, which leads to a reduced accuracy of translation start site prediction (Hyatt et al. 2010; Marcellin et al. 2013). To demonstrate that our approach can work beyond Bhen (38.2% GC), we have applied an earlier prototype on genomes with higher GC content: We could identify novel ORFs in the genome of *Burkholderia kirkii* (62.9% GC) including metabolic enzymes missed in a RAST annotation (Aziz et al. 2008), which carry out critical functions in the obligate symbiosis with plants of the genus *Psychotria* (Carlier et al. 2013). For *Bradyrhizobium diazoefficiens* (64.1% GC) and in combination with dRNA-seq data, we uncovered many novel short ORFs and internal start sites expressed under free-living conditions and in symbiosis with soybean (Cuklina et al. 2016). Finally, we applied our approach on shotgun proteomics datasets from *Escherichia coli* K-12 BW25113 (50.3% GC) during exponential growth (Krug et al. 2013) or grown under multiple conditions (Schmidt et al. 2015) (see Suppl. Methods). Even for the well-annotated genome of the parental strain of the Keio knockout strain collection (Baba et al. 2006), we could identify evidence for novel ORFs. These included 6 pseudogenes with solid expression evidence, but also short *in silico ORFs*, including a highly conserved sORF of 57 aa (3 peptides, 9 PSMs) plus several novel start sites (Tables S8, S9).

To enable proteogenomics for a larger microbiology research community with access to proteomics core facilities, we provide both a set of precomputed iPtgxDBs for several key prokaryotic model organisms including founder strains of gene knockout collections, and a step by step protocol (Figure 6). This can enable groups with bioinformatics support to generate iPtgxDBs for many newly sequenced organisms, e.g. type strains for the 11,000 named species targeted by the genomic encyclopedia of bacteria and archea (GEBA) project (Kyrpides et al. 2014) or environmental isolates. These organisms offer unique opportunities to study fundamental aspects such as the presence of novel biochemical reactions (Montes Vidal et al. 2017) or pathways relevant for biotechnological applications, the development and spread of antibiotics resistance (ABR), and key functionalities of important strains isolated from complex microbiomes, whose importance for e.g. human health (Cho and Blaser 2012) and plant protection from pathogen attack in agricultural settings (Berendsen et al. 2012), has been recognized.

**Figure 6.**
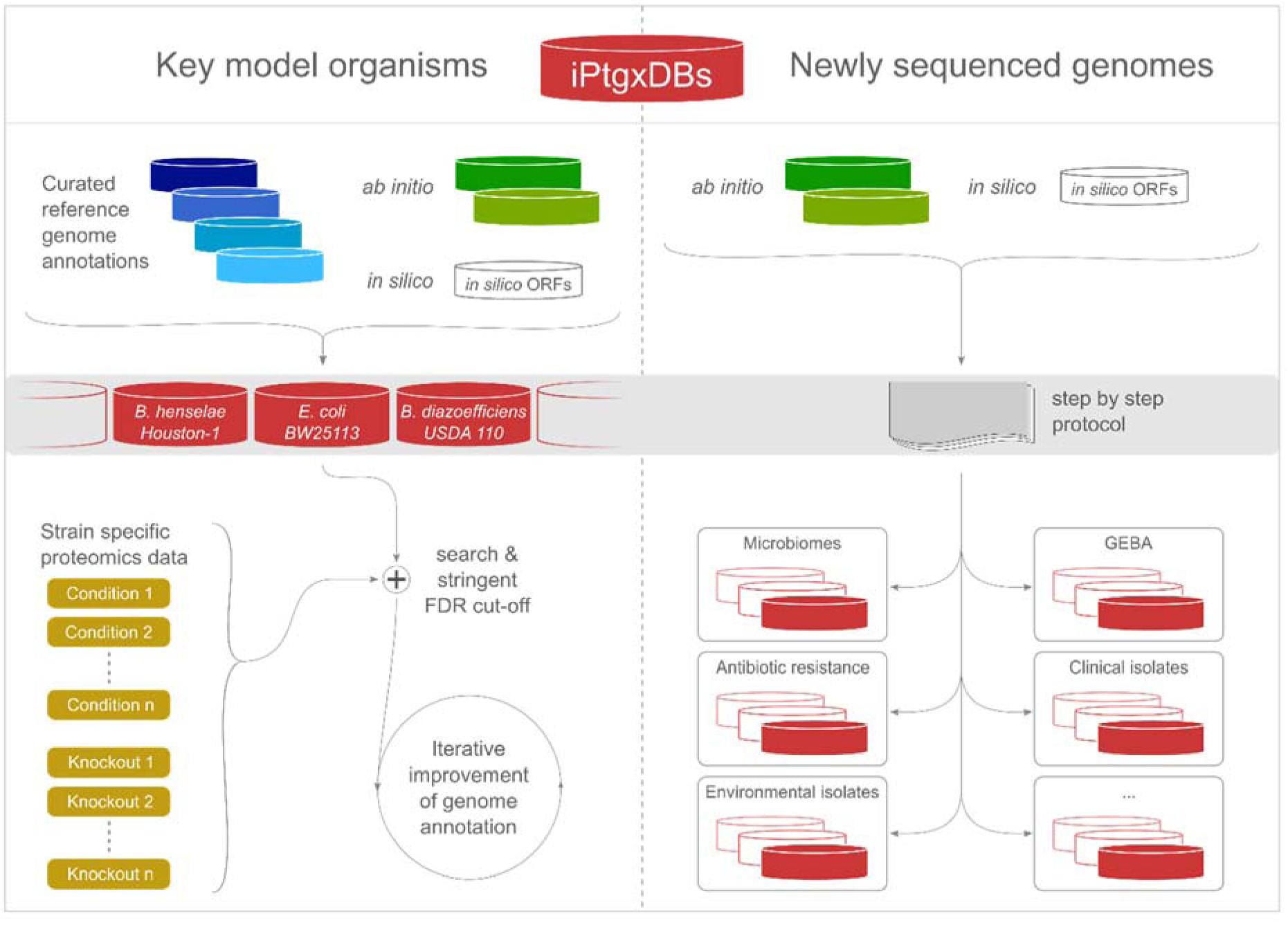
Application of our integrated proteogenomics approach. We release open source iPtgxDBs for several model organisms (https://iptgxdb.expasy.org), here for Bhen, *E. coli* BW25113 and *B. diazoefficiens* USDA 110 (left panel). Using proteomics data from any condition or knockout strain (light brown boxes, here schematically shown for *E. coli*), researchers can identify novelties, and iteratively improve the genome annotation e.g. in a community-driven genome Wiki approach (Salzberg 2007). The release of the software to integrate *ab initio* predictor(s) and *in silico* predictions (File S10) can help to improve genome annotations of many newly sequenced genomes (right panel).

## Discussion

We present a flexible, yet general proteogenomics strategy that allowed us to identify novelties in the genome of prokaryotes of different taxonomic origin (α-, β-, γ-proteobacteria) and widely ranging GC content. Investing a major effort in a pre-processing step to hierarchically integrate reference genome annotations and predictions into an iPtgxDB that covers the entire protein-coding potential pays off: close to 95% of the peptides unambiguously imply one protein (based on an extension of the PeptideClassifier concept (Qeli and Ahrens 2010) for prokaryotes), facilitating swift data analysis and mining. On top, informative identifiers capture overlap and differences of all resources, start codon and genomic coordinate information, such that novel ORFs, start sites or expressed pseudogenes can readily be identified and visualized. These features are unique to our solution. Our iPtgxDBs come in form of a protein search DB and a GFF file containing all annotations and identifiers.

For prokaryotes, the complexity of iPtgxDBs is lower than that of a regular protein search DB for e.g. yeast or human (Table S2). In our view, the benefit of generating a single iPtgxDB against which proteomics data from any condition (or knockout strain) can be searched to identify novel ORFs outweigh that of other elegant solutions that were developed for the more complex eukaryotes. Both splice graphs (Woo et al. 2013) and RNA-seq data (Wang et al. 2012) reduce the complexity and size of the search DB. However, in both cases, DBs specific for the conditions studied are generated, requiring bioinformatics expertise and limiting the general applicability of the resource. While our approach is unique, the GFF file can be very valuable for other proteogenomics software solutions like Genosuite (Kumar et al. 2013), PGP (Tovchigrechko et al. 2014) and PG Nexus (Pang et al. 2014), which allow users to search their data against a six-frame translation and later visualize identified peptides onto a genome sequence, but lack integrated and consolidated annotations.

The proteogenomics community is still to agree upon the best practice for required FDR thresholds and confirmation of novel candidates. Using very stringent FDR thresholds, as also advocated by Venter and colleagues (Venter et al. 2011), we show that the E-value distributions of PSMs for novel hits and target proteins are similar. Furthermore, we invested an extra effort to confirm the expression of novel ORFs with selective and sensitive PRM assays. The validation success (80% overall) ranged from 100% for SAAVs and highly expressed novel sORFs to around 55% for novel start sites. Reasons for the lower success with start sites can include N-terminal cleavage or modification, both of which can prevent detection of the single peptide to be confirmed (Goetze et al. 2009). Identification of internal start sites is even more difficult, but greatly benefits from the availability of dRNA-seq data (Cuklina et al. 2016) and/or N-terminal enrichment steps.

Focusing on the description of our novel strategy, we did not further characterize or functionally validate novel sORFs beyond the PRM confirmation. More effort will be required to assess the functional relevance of sORFs, e.g. by individual gene deletion or genome-wide transposon mutagenesis screens (Christen et al. 2011). Recent work in yeast suggested that sORFs may represent a pool of proto-genes that are under evolutionary pressure and may lead to the birth of novel genes (Carvunis et al. 2012). Indeed, genes that emerged more recently tend to be shorter (Tautz and Domazet-Loso 2011).

Besides identifying missed sORFs, our data indicated that i) the procedures to annotate pseudogenes differ between resources (and even releases) and have to be treated with caution, and ii) likely over-predicted ORFs can be uncovered by relying on complete, condition-specific expressed proteomes. Recent advances to comprehensively identify expressed proteomes within a few days (Nagaraj et al. 2012; Richards et al. 2015) suggest that proteomics data can, at least in part, address this issue of over-prediction. Such extensive datasets can also uncover functionally relevant genomic changes down to the SAAV level, with implications for clinical proteomics and beyond. For example, by tracking clinically relevant pathogens either over time (Lee et al. 2017) or comparing different strains, genome changes that correlate with higher pathogenicity (de Souza et al. 2011; Nasser et al. 2014; Malmstrom et al. 2015) can be identified, some of which ideally are supported by direct protein expression evidence for SAAVs.

Importantly, our data show that assembling the correct genome sequence of the strain under study is of critical importance: it is the optimal basis not only to comprehensively identify expression differences between the conditions studied, but also to accurately identify novel sORFs by proteogenomics. An initial *de novo* assembly should thus be carried out routinely for the most important strains, in particular those that form the basis for long-term projects aiming to integrate functional genomics data.

We favor a conservative approach to genome re-annotation, ideally carried out by consortia that iteratively improve the annotation of their respective model or non-model organisms (Armengaud et al. 2014), e.g. relying on a genome Wiki concept (Salzberg 2007) (Figure 6). By releasing iPtgxDBs initially for three model organisms (https://iptgxdb.expasy.org) and the software to create them (File S10), we hope to enable a large user base to apply proteogenomics in the initial genome annotation step. This will provide an optimal basis for systems-wide functional studies and genome-scale regulatory or metabolic predictions, and help to fully capitalize on the genome information and decode its function.

## Data access

All data are publicly available: the genome sequence of Bhen variant-1 strain CHDE101 (Genbank; acc. # CP020742), the PRM data (Email: panorama+wollscheid@p rote in ms. net; Password:!9QcZ4#T) under https://panoramaweb.org/labkey/Bartonella_Proteogenomics.url, and iPtgxDBs for Bhen Houston-1 (RefSeq NC_005956), Bhen CHDE101, *E. coli* BW25113 and *B. diazoefficiens* USDA 110 (in both FASTA format and as GFF files to visualize all integrated annotations); they can be downloaded from https://iptgxdb.expasy.org (user: preview, pw: iptgxdblive). Scripts to generate iPtgxDBs for any prokaryote are available in the Supplement (File S10).

### Acknowledgements

We thank A. Wicki for initial work on the genome assemblies, R. Schlapbach for access to the FGCZ and continued support, F. Freimoser, G. Pessi and 3 anonymous reviewers for valuable input on the manuscript, and sciCORE for hosting the iPtgxDB website. The project was supported by BW through the D-HEST BioMedical Proteomics Platform (BMPP). CHA acknowledges funding for UO and ARV from the SNSF under grant 31003A-156320.

## Author contributions

UO and CHA devised the integrated proteogenomics strategy; UO, ARV and CHA analyzed data; UO, ARV and MS made figures; ARV, MS and DM assembled and annotated the genomes with input from AP, JEF, ON and MR; MQ and CD provided Bhen gDNA and new protein extracts; SG and BW validated selected candidates by PRM, MB developed the iPtgxDB website with input from ARV and CHA; CHA wrote the manuscript. All authors commented on the manuscript.

## Methods

### Source of reference genome annotations and *ab initio* predictors

Annotations of the Bhen reference genome (Alsmark et al. 2004) were obtained from NCBI’s RefSeq (Pruitt et al. 2012) (NC_005956.1; from 06/10/2013 called RefSeq2013, and 07/30/2015, called RefSeq2015), from Ensembl’s Genomes project (GCA_000046705.1, Feb/2015), and from Genoscope’s microbial genome annotation & analysis platform (Vallenet et al. 2013) (v2.7.3, accessed 03/09/2016). *Ab initio* gene predictions from Prodigal (Hyatt et al. 2010) (v2.6) and ChemGenome (Singhal et al. 2008) were used (v2.0, http://www.scfbio-iitd.res.in/chemgenome/chemgenomenew.jsp; with parameters: method, Swissprot space; length threshold, 70 nt; initiation codons, ATG, CTG, TTG, GTG). Files were parsed to extract the identifier, coordinates and sequences of *bona fide* protein-coding sequences (CDS) and pseudogene entries.

### Integrative proteogenomics approach

The annotations were collapsed into singletons (same sequence in all sources) or annotation clusters of two or more sequences with the same stop codon but different start sites. For clusters, we define an anchor sequence from the annotation highest up in the hierarchy, e.g. RefSeq2015. We construct an informative and transparent protein identifier that integrates all relevant information: a code is added to the anchor sequence for each identical annotation (RefSeq2013=rso, Ensembl=ens, Genoscope=geno, Prodigal=prod, ChemGenome=chemg, *in silico* ORF=orf) separated by a pipe sign (e.g. BH_RS00220|rso|ens|geno). Identical *in silico* ORFs are not considered. For alternative start sites, the length difference compared to the anchor annotation is added prior to the code (e.g. …|-17aa_prod|+6aa_chemg). Finally, chromosome, start and stop position, reading frame, start codon and CDS length complete the identifier. The anchor sequence identifier thus integrates relevant information of the genomic location and all annotation sources for this region, including possible reductions and extensions (see Figure 2A). Identifiers for entries with alternative initiation sites contain a reference to the anchor annotation, the length difference and the annotation source (e.g. BH_RS00220_+6aa_chemg). To create an iPtgxDB, we only add the complete sequence of the anchor sequence of a cluster plus sequences of N-terminal regions that give rise to identifiable (i.e. different from the anchor sequence) tryptic peptides for the additional proteoforms of the cluster. For extensions, the N-terminal sequence up to the first tryptic cleavage site in the anchor sequence is added, for internal start sites the N-terminal tryptic peptide if it starts from an alternative initiation codon other than ATG (TTG, GTG, or CTG) giving rise to a N-terminal Met instead of a Leu, Val, or Leu, respectively. For pseudogenes, we added the suffix “_p” (e.g. BH_RS02905_p) or “_fCDS_p” (e.g. BHGENO0333_fCDS_p; “fragmented CDS”, for Genoscope pseudogenes) to the identifier, and a sequence translated to the first stop codon to the protein DB. *In silico* ORFs above a selectable length threshold (18 aa) were added (Suppl. Methods). For the *de novo* assembled Bhen MQB277 genome, Prodigal predictions and *in silico* ORFs were integrated (same length cut-off).

### PeptideClassifier analysis of protein search DBs

The complexity and redundancy of protein search DBs was assessed with the web-based PeptideClassifier tool (http://peptideclassifier.expasy.org) to derive an evidence class for every tryptic peptide of 6 to 40 aa (Figure S2). To deal with multiple different annotations for the same gene model, we generated the required gene-annotation mapping files using the stop codon coordinates as common gene name across annotations, i.e. an extension of the original concept of gene-protein mapping (Qeli and Ahrens 2010) (see text and Figure 2). A webservice to support peptide classification for proteogenomics in prokaryotes will soon be released.

### Stringent re-analysis of proteomics and transcriptomics data

Proteomics data (ProteomeXchange, acc.# PXD000153) was searched with MS-GF+ (v.10.0.72) (Kim and Pevzner 2014) and described parameters (Omasits et al. 2013) against the RefSeq2015-based DB, the iPtgxDB (51,541 proteins), and the iPtgxDB of our *de novo* assembly (52,687 proteins). A PSM FDR threshold of 0.01% was used, estimated peptide and protein level FDRs were 0.12% and 0.6%, respectively (Table S7). Protein expression estimates and differential expression values were computed as described (Omasits et al. 2013). Reads from the matched Bhen transcriptomics dataset (GEO, acc.# GSE44564) were stringently re-mapped both to the NCBI reference genome (Alsmark et al. 2004) and to our *de novo* assembly using NovoalignCS v1.06.04 (Novocraft, Selangor, Malaysia). For reads supporting coding SNVs, we only considered reads without a mismatch. This allowed us to provide transcriptomic support for a substantial amount of observed genomic differences, both coding and non-coding. For more details, see Suppl. Methods.

### Bacterial strains, genomic DNA and protein extracts

High quality gDNA was extracted from Bhen strains MQB277 and CHDE101 (Schmid et al. 2004), a close laboratory variant of the NCBI Bhen Houston-1 ATCC49882 reference strain (Alsmark et al. 2004) and parental strain of MQB277, using Sigma’s GenElute kit. Both were sequenced (PacBio) and assembled into one high quality contig (Suppl. Methods). Protein extracts of cytoplasmic (cyt) and total membrane (TM) fractions were prepared from bacterial cells grown under uninduced and induced conditions as described (Omasits et al. 2013).

### Protein features, functional annotation, conservation

Several protein features including signal peptides, transmembrane topology, lipoproteins, and protein domains were predicted, and protein sequences functionally annotated by EggNOG. Conservation of novel ORFs was assessed with tblastn. Predominant SCL information was computed for all proteins (including novel ORFs) similar to (Stekhoven et al. 2014). For details, see Suppl. Methods.

### Independent validation by targeted proteomics

Peptides for novel ORFs, start sites, expressed pseudogenes, or assembly-specific changes were selected based on spectral count, number of tryptic sites, number of missed cleavage sites, and PeptideRank prediction (Qeli et al. 2014). Heavy-labeled reference peptides were purchased from JPT Peptide Technologies GmbH (Berlin, Germany) and used to set up PRM assays (Peterson et al. 2014). Specific transitions were measured in cyt and TM extracts of biological replicates of both conditions (new fractions). Only traces within a mass accuracy of 10 ppm were evaluated; we excluded transition interference by manually validating co-elution of peptide traces. For details of the sample preparation and MS set-up see Suppl. Methods.

